# Heterogeneity of anti-Caspr2 antibodies: specificity and pathogenicity

**DOI:** 10.1101/2025.01.16.633238

**Authors:** Julia Su, Rohan Gupta, Scott Van Hoof, Jakob Kreye, Harald Prüss, Benjamin Spielman, Lior Brimberg, Bruce T. Volpe, Patricio T. Huerta, Betty Diamond

## Abstract

Maternal anti-Caspr2 (Contactin-associated protein-like 2) antibodies have been associated with increased risk for autism spectrum disorder (ASD). Previous studies have shown that *in utero* exposure to anti-Caspr2 antibodies results in a phenotype with ASD-like features in male mice. Here we ask whether four newly generated antibodies against Caspr2 are pathogenic to the developing fetal brain and whether they function through similar means. Our results show that the novel anti-Caspr2 antibodies recognize different epitopes of Caspr2. *In utero* exposure to these antibodies elicits differential ASD-like phenotypes in male offspring, tested in the social interaction, open field, and light-dark tasks. These results demonstrate variability in the antigenic specificity and pathogenicity of anti-Caspr2 antibodies which may have clinical implications.

## INTRODUCTION

Autism spectrum disorder (ASD) affects 1 in 36 children in the United States, with a four-fold higher prevalence in males [1]. This heterogeneous neurodevelopmental disorder is characterized by impaired social interaction, communication deficits, and restricted, repetitive patterns of behavior or activities [1]. ASD presents across a wide phenotypic spectrum, ranging from individuals with severe intellectual disability and seizures requiring lifelong care to those with milder presentations who live independently but experience challenges with social skills and communication [4]. While the etiology of ASD is complex and likely involves both genetic and environmental factors [5, 6], increasing evidence points to a role for maternal autoantibodies in some cases. Women with autoimmune diseases have an increased risk of having a child with ASD [7–9]. Studies have revealed that a significant percentage of mothers of children with ASD possess elevated levels of anti-brain antibodies compared to mothers of typically developing children [7, 10, 11]. These antibodies have been detected through Western blot analysis for IgG reactivity to brain lysate and immunohistology for reactivity to intact brain tissue, suggesting a potential link between maternal autoimmunity and ASD. Furthermore, animal models demonstrate that maternal anti-brain antibodies can cross the placenta and affect fetal neurodevelopment, resulting in offspring displaying ASD-like behaviors [12–14].

Contactin-associated protein-like 2 (Caspr2, encoded by the ASD risk gene *CNTNAP2*) has been identified as a target of some of these maternal anti-brain antibodies [13, 15]. Caspr2, a cell adhesion protein of the neurexin family, is crucial for stabilizing the voltage-gated potassium channel complex at the juxtaparanode on the myelinated axon in the postnatal brain [19]. However, during development, Caspr2 is thought to play a role in neuronal migration and synapse stabilization [18, 20, 21]. Our previous work demonstrated that 37% of mothers with brain-reactive antibodies and a child with ASD harbor anti-Caspr2 antibodies [13]. Interestingly, Caspr2 knockout (KO) mice exhibit an ASD-like behavioral phenotype [16– 18], further implicating Caspr2 dysfunction in ASD pathogenesis.

We have previously isolated a monoclonal anti-Caspr2 IgG antibody from a mother with brain-reactive antibodies and a child with ASD. Prenatal exposure to this antibody in mice resulted in male-specific alterations in cortical development, decreased dendritic arborization in pyramidal neurons within the CA1 region of the hippocampus, and behavioral changes reminiscent of ASD, including impaired sociability, flexible learning deficits, and repetitive behaviors [13]. A subsequent study using a polyclonal antibody approach, where female mice were immunized with the extracellular portion of Caspr2, yielded similar developmental and behavioral alterations in male offspring [15]. These findings strongly support the hypothesis that maternal anti-Caspr2 antibodies can disrupt neurodevelopment, particularly in male offspring [13]. The present study investigates a panel of monoclonal anti-Caspr2 antibodies to determine their pathogenicity, binding specificity, and associated clinical phenotypes.

## RESULTS

### Generation of anti-Caspr2 antibodies

Four unique monoclonal anti-Caspr2 antibodies were generated via hybridoma technology from spleen cells of female mice immunized with the extracellular domain of human Caspr2. Two IgG1 antibodies, termed P9C7 and P11G7, were derived using the P3-X63-Ag8 myeloma line while two IgG2b antibodies, termed P6B2 and P13F4, were generated using the NSO-BCL2 (Bcl-2 overexpressing) myeloma line.

### Characterization of IgG1 anti-Caspr2 antibodies, P11G7 and P9C7

Sequence analysis confirmed the unique heavy and light chain sequences of P11G7 and P9C7 (**Fig. 1A**). Moreover, ELISA analysis showed that both molecules were IgG1 antibodies (**Fig. 1B**). Importantly, P11G7 and P9C7 displayed robust binding to human and mouse Caspr2 expressed in HEK293 cells, while an IgG1-matched isotype control (CON1) showed null binding (**Fig. 1C**, *top*). P11G7 and P9C7 also recognized human and mouse Caspr2 in GnTI-HEK293 cells (lacking complex glycosylation) (**Fig. 1C**, *bottom*). In mouse brain tissue, P11G7 bound to PFA-fixed adult, snap-frozen adult, and PFA-fixed E15.5 embryonic brain sections (**Fig. 1D**, *top*). P9C7 exhibited binding to snap-frozen adult and E15.5 brain sections (**Fig. 1D**, *bottom*). Both antibodies also bound to Caspr2 in primary neuronal cultures, colocalizing with a commercial anti-Caspr2 antibody (**Fig. 1E**).

**Figure 1:**
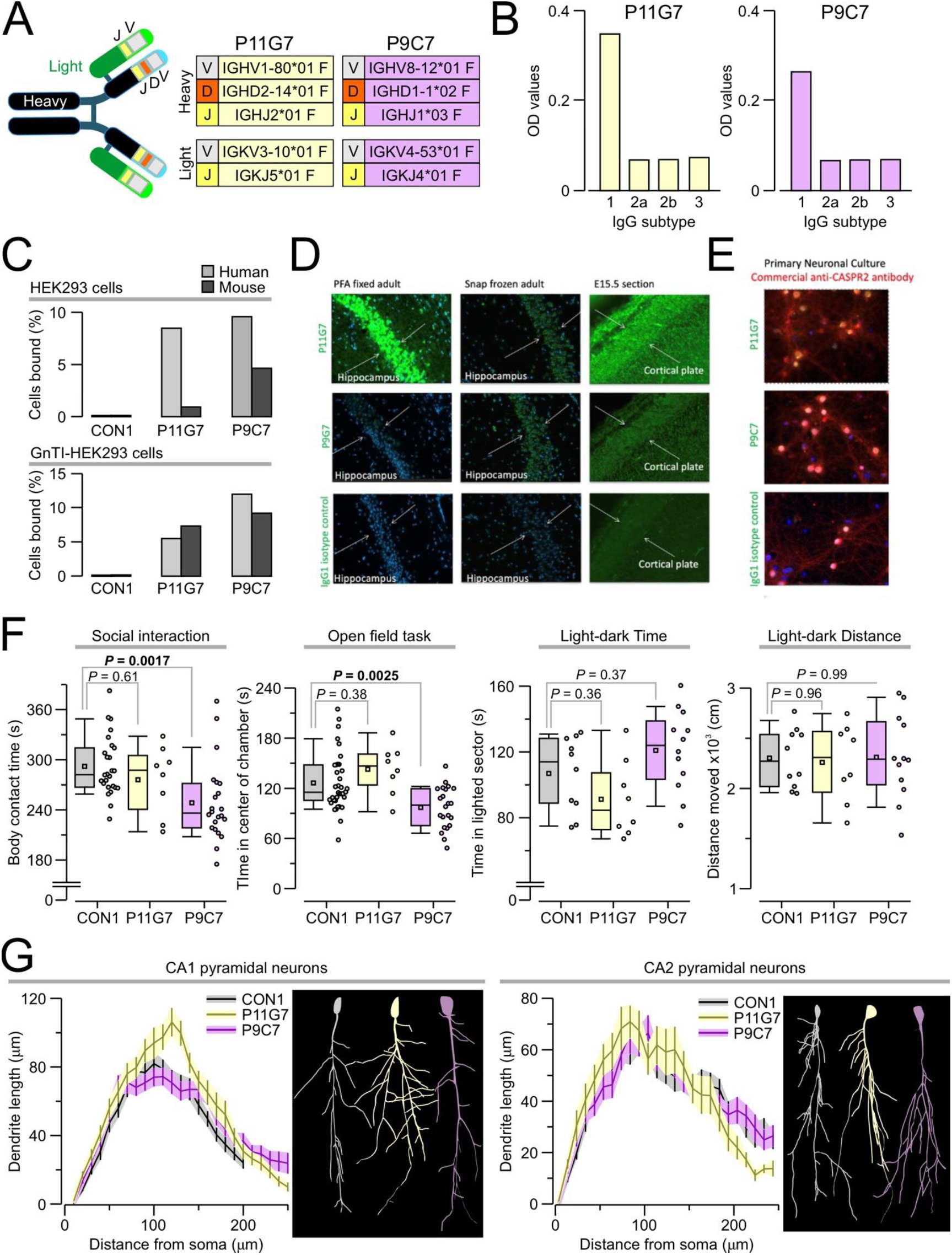
IgG1 anti-Caspr2 antibodies. **(A)** Diagram of the structure of P11G7 and P9C7 with their unique sequences for the heavy and light chains. **(B)** Isotype determination with a sandwich ELISA done by coating a plate with various anti-IgG isotype subtypes, adding purified antibody to each well, and measuring their optical density (OD). P11G7 and P9C7 are clearly IgG1 isotype. **(C)** Anti-Casp2 binding on HEK293 cells (top) and GnTI-HEK293 cells (bottom). P11G7 and P9C7 bind to both human Casp2 and mouse Caspr2 compared to an IgG1-matched isotype control (CON1). **(D)** Binding of P11G7 (10 μg/mL) and P9C7 (10 μg/mL) to adult mouse brain sections (PFA-fixed or snap-frozen) and E15.5 whole embryo sections. P11G7 binds to all sections, whereas P9C7 binds to snap-frozen and embryonic sections but not PFA-fixed sections. Representative images of the CA1 region of the hippocampus (adult) and cortical plate (E15.5) at 20x. **(E)** Binding of P11G7 (10 μg/mL) and P9C7 (10 μg/mL) to primary neuronal cultures (fixed at DIV14) stained with primary antibodies. Hybridoma anti-Caspr2 antibodies are labeled in green, a commercial anti-Caspr2 antibody (ABN1380) in red, and their colocalization in yellow. Images taken at 40x. **(F)** Assessment of male mice exposed to anti-Caspr2 antibodies *in utero* reveals that mice in the P11G7 group have no differences in any of the tasks compared to the CON1 group, while mice in the P9C7 group has less body contact with an unfamiliar mouse in the social interaction task, and less time in the center of the chamber in the open field task. Statistical testing done with ANOVA followed by Tukey post-hoc test. **(G)** Sholl analysis of pyramidal neurons from the CA1 and CA2 regions of the hippocampus reveals no significant differences in dendritic length between the groups. Insets (at right in each panel) show representative tracing of CA1 and CA3 neurons.

The model of prenatal exposure to maternal antibodies was used in three groups of offspring: those from dams exposed to P11G7, dams exposed to P9C7, and dams exposed to CON1. At a baseline level, no significant differences in body weight, coat, grip strength, body tone, reflexes, or sickness behavior were observed in male or female offspring exposed to any of the anti-Caspr2 antibodies compared to isotype controls. Moreover, female offspring of all groups showed no significant differences in the social interaction, open field, and light-dark tasks (data not shown). Importantly, P9C7 male offspring displayed reduced time interacting in the social task (**Fig. 1F**, mean ± SEM, CON1 = 292.01 ± 6.98 s; P9C7 = 248.49 ± 10.04 s; *q* = 5.15, *P* = 0.0017, ANOVA with Tukey test) and increased anxiety-like behavior in the open field task (**Fig. 1F**, time in center, CON1 = 126.43 ± 5.73 s; P9C7 = 97.03 ± 5.37 s; *q* = 4.95, *P* = 0.0025). Conversely, *in utero* exposure to P11G7 did not result in significant changes during social interaction (**Fig. 1F**, P11G7 = 276.04 ± 14.6 s; *q* = 1.36, *P* = 0.61) and open field behavior (**Fig. 1F**, P11G7 = 142.71 ± 10.48 s; *q* = 1.89, *P* = 0.38) in male offspring. Additionally, no differences were observed in the in the light-dark task for time spent in the lighted sector (**Fig. 1F**, CON1 = 106.91 ± 7.17 s; P9C7 = 120.96 ± 7.33 s; *q* = 1.94, *P* = 0.37; P11G7 = 91.19 ± 8.29 s; *q* = 1.95, *P* = 0.36) and the distance moved (**Fig. 1F**, CON1 = 2302.24 ± 93.86 cm; P9C7 = 2310.23 ± 124.06 cm; *q* = 0.07, *P* = 0.99; P11G7 = 2259.37 ± 138.06 cm; *q* = 0.34, *P* = 0.96) for any of the antibody-exposed groups.

A neuroanatomical approach, with Golgi-Cox staining followed by Sholl analysis, was applied to quantify the dendritic arborization of pyramidal neurons in the CA1 and CA2 regions of the hippocampus. Surprisingly, no significant changes in dendritic length were observed in the CA1 or CA2 hippocampal regions of P9C7-exposed males (**Fig. 1G**). The control values for dendritic length were comparable to values in animals of similar age (12–14 months) that we have reported [46]. Similarly, *in utero* exposure to P11G7 did not result in significant anatomical changes in CA1 and CA2 neurons of male offspring (**Fig. 1G**).

### Characterization of IgG2b anti-Caspr2 antibodies, P13F4 and P6B2

Heavy and light chain sequences were obtained for P13F4 and P6B2 (**Fig. 2A**). Moreover, ELISA analysis showed that both molecules were IgG2 antibodies (**Fig. 2B**). Moreover, P13F4 bound more robustly to human than mouse Caspr2 in both HEK293 and GnTI-HEK293 cells (**Fig. 2C**), suggesting a stronger binding selectivity for human Caspr2. P6B2 showed a similar pattern, binding better to human Caspr2 than mouse Caspr2 in HEK294 and GnTI-HEK293 cells (**Fig. 2C**). Importantly, an IgG2-matched isotype negative control (CON2) showed null binding to HEK293 and GnTI-HEK293 cells (**Fig. 2C)**. In brain tissue, P6B2 bound only to PFA-fixed adult sections, whereas P13F4 bound to both PFA-fixed adult and E15.5 embryonic sections (**Fig. 2D**). Both P6B2 and P13F4 bound to Caspr2 in neuronal cultures (**Fig. 2E**).

**Figure 2:**
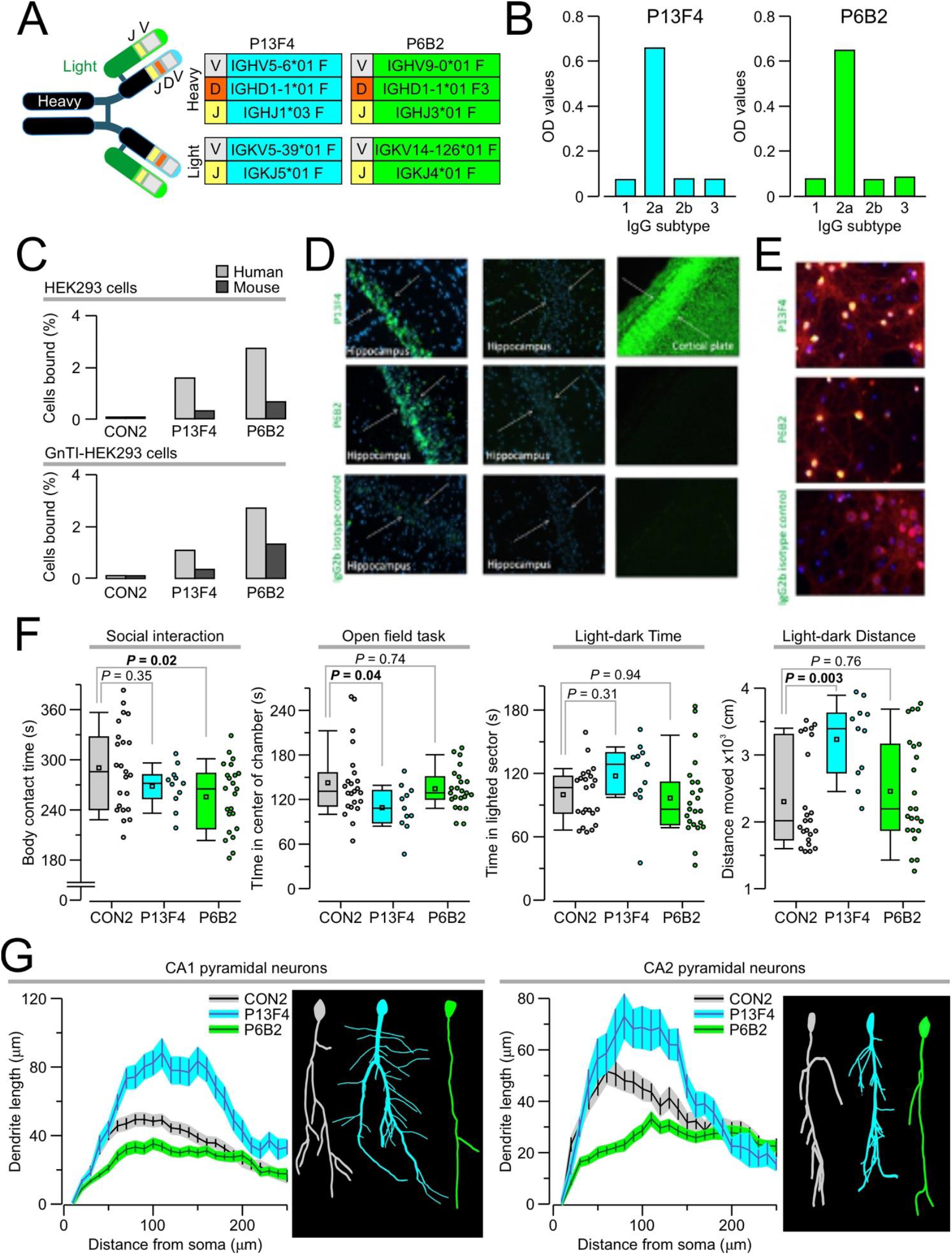
IgG2 anti-Caspr2 antibodies. **(A)** Diagram of the structure of P13F4 and P6B2 with their unique sequences for the heavy and light chains. **(B)** Isotype determination with a sandwich ELISA done by coating a plate with various anti-IgG isotype subtypes, adding purified antibody to each well, and measuring their optical density (OD). P13F4 and P6B2 are IgG2 isotype. **(C)** Anti-Casp2 binding on HEK293 cells (top) and GnTI-HEK293 cells (bottom). P13F4 and P6B2 bind preferentially to human Casp2 and less to mouse Caspr2 compared to the null binding of an IgG2-matched isotype control (CON2). **(D)** Binding of P13F4 (10 μg/mL) and P6B2 (10 μg/mL) to adult mouse brain sections (PFA-fixed or snap-frozen) and E15.5 whole embryo sections. In brain tissue, P6B2 binds only to PFA-fixed adult sections, whereas P13F4 binds to both PFA-fixed adult and E15.5 embryonic sections. Representative images of the CA1 region of the hippocampus (adult) and cortical plate (E15.5) at 20x. **(E)** Binding of P13F4 (10 μg/mL) and P6B2 (10 μg/mL) to primary neuronal cultures (fixed at DIV14) stained with primary antibodies. Hybridoma anti-Caspr2 antibodies are labeled in green, a commercial anti-Caspr2 antibody (ABN1380) in red, and their co-localization in yellow. Images taken at 40x. **(F)** Assessment of male mice exposed to anti-Caspr2 antibodies *in utero* reveals that mice in the P13F4 group have normal behavior in the social interaction task but less time in the center of the chamber during the open field task and more distance covered in the light-dark box task. Mice in the P6B2 group show less body contact with an unfamiliar mouse in the social interaction task. Statistical testing done with ANOVA followed by Tukey post-hoc test. **(G)** Sholl analysis of pyramidal neurons from the CA1 and CA2 regions of the hippocampus reveals that P13F4-exposed males have a significant increase in dendritic length for CA1 (*P* = 0.0001) and CA2 (*P* = 0.0069) neurons, whereas P6B2-exposed males have significantly shorter dendrite length when compared to male mice exposed to isotype control in both CA1(*P* < 0.0001) and CA2 neurons (*P* = 0.0001). Insets (at right in each panel) show representative tracing of CA1 and CA3 neurons.

Male offspring exposed *in utero* to P13F4 exhibited normal time interacting in the social interaction task (**Fig. 2F**, CON2 = 290.21 ± 10.6 s; P13F4 = 268.19 ± 7.65 s; *q* = 1.98, *P* = 0.35, ANOVA with Tukey test) but increased anxiety-like behavior in the open field task (**Fig. 2F**, CON2 = 142.68 ± 10.6 s; P13F4 = 109.04 ± 9.27 s; *q* = 3.43, *P* = 0.04) and hyperactivity in the light-dark box task in terms of distance moved (**Fig. 2F**, CON2 = 2302.42 ± 152.91 cm; P13F4 = 3233.42 ± 177.73 cm; *q* = 4.88, *P* = 0.003). Moreover, P13F4-exposed males showed a significant increase in dendritic length for CA1 and CA2 neurons (**Fig. 2G**).Remarkably, P6B2-exposed males displayed reduced social interaction (**Fig. 2F**, P6B2 = 255.58 ± 8.43 s; *q* = 3.87, *P* = 0.022) and no changes in anxiety-like behavior in the open field task (**Fig. 2F**, P6B2 = 134.48 ± 5.75 s; *q* = 1.04, *P* = 0.74) and the light-dark task (Fig. 2F, P6B2 = 2457.62 ± 165.76 cm; *q* = 1.01, *P* = 0.76). Finally, P6B2-exposed males had significantly decreased dendritic length within CA1 and CA2 neurons (**Fig. 2G**).

### Anti-Caspr2 antibodies bind to different epitopes on Caspr2

To determine the target domain recognized by each anti-Caspr2 antibody, HEK293T cells were transfected with either the full length or a mutant construct encoding for the human Caspr2 protein. Each mutant construct contained the deletion of one of the eight extracellular domains or had only a single domain (**Fig. 3A**). These epitope mapping studies using human Caspr2 deletion mutants revealed that both P11G7 and P9C7 targeted the discoidin domain (**Fig. 3B**), as they bound to every mutant Caspr2 construct except the construct lacking the discoidin domain (ΔDisc). Also, both P11G7 and P9C7 bound to a construct containing only the discoidin domain (**Fig. 3C**).

**Figure 3:**
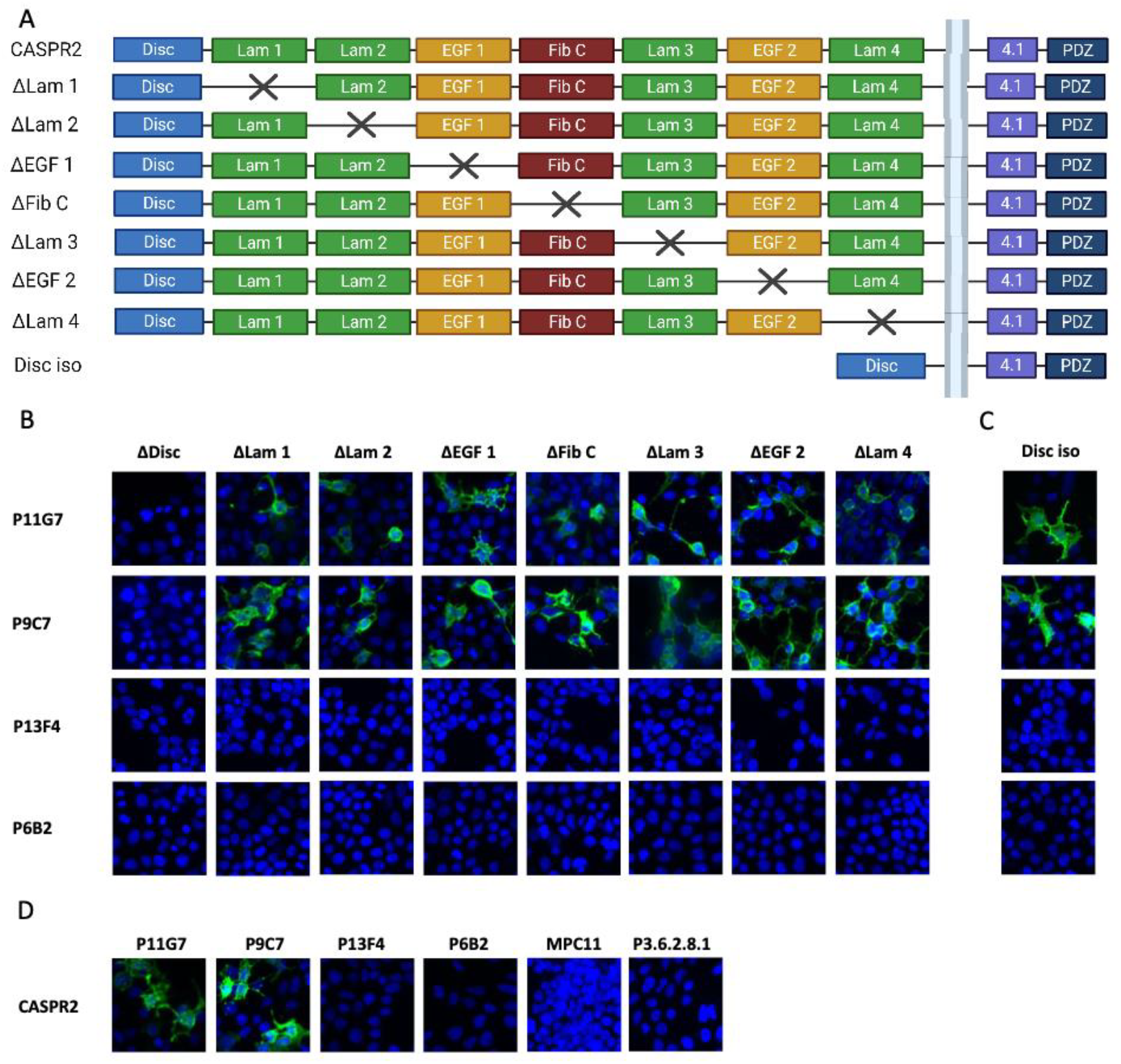
Identification of anti-Caspr2 antibody target domains. **(A)** CASPR2 mutant constructs. The extracellular portion of CASPR2 has 8 domains. The mutant constructs each missing one of the domains were previously generated (van Hoof et al.). The Disc iso construct contains only the discoidin domain. **(B)** HEK293T cells were transfected with one of the CASPR2 mutant constructs for 48 h and incubated with each anti-CASPR2 antibody using live staining. P11G7 and P9C7 bind to all mutant constructs except the one missing the discoidin domain. P6B2 and P13F4 do not bind to any of the constructs. Isotype controls showed no binding to any construct. **(C)** P11G7 and P9C7 bind to the discoidin domain only construct. P13F4 and P6B2 do not bind to the discoidin domain only construct. **(D)** P11G7, P9C7, bind to CASPR2 transfected HEK293T cells. P6B2, P13F4 and the isotype controls do not.

In contrast, P6B2 and P13F4 did not bind to any of the deletion mutants. (**Fig. 3B-C**), consistent with the preferential binding of P6B2 to mouse Caspr2 and preferential binding of P13F4 to Caspr2 in the absence of complex glycosylation.

## DISCUSSION

Our previous work identified anti-Caspr2 IgG in a substantial proportion (36%) of mothers with anti-brain antibodies and a child with ASD, suggesting Caspr2 as a potential target in ASD pathogenesis [7]. Given the prevalence of these antibodies, we hypothesized that multiple pathogenic mechanisms for Caspr2 antibodies might exist. To investigate this proposition, we generated and characterized a panel of monoclonal anti-Caspr2 antibodies. Our findings reveal distinct differences in their specificity and pathogenicity, contributing to the heterogeneous nature of ASD.

## IgG1 antibodies (P11G7 and P9C7) have divergent effects despite shared target domain

Both P11G7 and P9C7 target the discoidin domain of Caspr2, similar to Caspr2 autoantibodies from autoimmune encephalitis patients that were found to predominantly bind the discoidin domain [42– 44]. Despite this shared specificity, they induced distinct behavioral outcomes in male offspring following *in utero* exposure. While P11G7 exposure resulted in no overt behavioral changes, P9C7-exposed males exhibited both social interaction deficits and increased anxiety-like behavior. This divergence suggests that even antibodies targeting the same domain can differentially impact neurodevelopment, possibly through subtle variations in binding affinity, different epitopes within the same target domain, post-translational modifications of the Fc part, downstream signaling pathways, or by influencing distinct neuronal populations. The lack of dendritic changes in the CA1 and CA2 pyramidal neurons of P9C7-exposed males suggests that its effects on social behavior may involve other brain regions or processes.

## IgG2b antibodies (P6B2 and P13F4) show distinct binding profiles and behavioral outcomes

P6B2 and P13F4 each exhibited distinct binding characteristics. P6B2 preferentially bound mouse Caspr2; P13F4showed binding to both human and mouse Caspr2 only in the absence of complex glycosylation, suggesting its epitope is masked in the native protein. *In utero* exposure to P6B2 resulted in social deficits in male offspring, accompanied by a significant reduction in dendritic length in the CA1 and CA2 pyramidal neurons, providing a potential anatomical correlate for the observed behavioral changes. In contrast, P13F4 exposure led to hyperactivity and increased anxiety-like behavior, along with increased dendritic length in the same hippocampal regions. This contrasting effect on dendritic morphology highlights the diverse ways in which anti-Caspr2 antibodies can disrupt neurodevelopment.

## Potential mechanisms and the male bias in ASD

The distinct behavioral and anatomical changes induced by these antibodies suggest multiple pathways through which anti-Caspr2 antibodies might contribute to ASD-like phenotypes. The observed social deficits, particularly with P6B2 exposure, align with previous studies demonstrating reduced dendritic arborization in the hippocampus following *in utero* exposure to anti-Caspr2 antibodies [13, 15]. The hyperactivity observed in P13F4-exposed males, coupled with increased dendritic length, warrants further investigation into the specific neuronal circuits involved. While previous studies using homocysteine models have shown similar hyperactivity and dendritic changes [31], the distinct molecular targets suggest different mechanisms may be at play in our anti-Caspr2 model.

The male-specific vulnerability to anti-Caspr2 antibodies observed in our study is consistent with the male bias in ASD prevalence and previous findings in anti-Caspr2 models [13, 15]. This suggests a potential role for sex chromosomes in mediating this differential susceptibility [38], although the precise mechanisms remain to be elucidated.

## Conclusion

Our findings demonstrate that monoclonal anti-Caspr2 antibodies, even those targeting the same domain, can induce a range of ASD-like phenotypes in mice, highlighting the complexity of ASD pathogenesis. Further investigation into the specific molecular mechanisms underlying these diverse effects will be crucial for understanding the heterogeneity of ASD and developing targeted therapeutic strategies.

## METHODS

### Mice and study approval

C57BL/6 mice were purchased from The Jackson Laboratory. Studies were carried out in strict accordance with the Guide for the Care and Use of Laboratory Animals of the NIH. The protocol was approved by the Institutional Animal Care and Use Committee of the Feinstein Institutes for Medical Research.

### Hybridomas secreting anti-Caspr2 antibodies

C57BL/6J mice (6–8-week-old females) were immunized with 200 μg of the extracellular portion of human Caspr2 (emulsified 1:1 in Complete Freund’s Adjuvant) via intraperitoneal injection.Mice were boosted 3 weeks after the initial immunization with 200 μg of Caspr2 (emulsified 1:1 in Incomplete Freund’s Adjuvant), via intraperitoneal injection. To confirm that female mice generated an immune response to Caspr2, antibody titers were checked via HEK Cell Based Assay as described in [Brimberg; Bagnall], Three days after the booster, mice were euthanized and spleens were harvested for splenocytes. Splenocytes were gently extracted using the balloon technique. Briefly, the spleen was placed in a dish with DMEM (with high glucose and 1% penicillin and streptomycin), threaded lengthwise along a 22-gauge needle syringe containing DMEM, which was gently expelled from the syringe to push out splenocytes in a single cell suspension, which was pipetted gently up and down. Splenocytes were transferred to a 50-mL conical tube and placed on ice for 5 min to allow debris to separate from cells by sedimentation. After sedimentation, the top portion of the splenocyte mixture was transferred to a clean 50-mL conical tube. To remove red blood cells from the splenocyte mixture, the mixture was pelleted by centrifugation at 400 g for 5–10 min. After the supernatant was removed, the pellet was loosened by flicking, and 5 mL of cold NH4Cl (0.01 mL, 7M pH 7) was added. The tube was placed on ice and incubated for 10 min (with a brief tube swirl at the 5-min mark). At 10 min, 10 mL of cold DMEM was added slowly while swirling the tube to stop the lysis reaction. Once 10 mL of DMEM had been added, the remainder of the tube was filled with cold DMEM, pelleted by centrifugation, and resuspended in 10 mL of fresh DMEM to fully remove the lysis buffer. Splenocytes were combined with myeloma cells. Myeloma cells were grown separately in preparation for fusion with splenocytes (myeloma lines were obtained from Dr. Matthew Scharff, Albert Einstein College of Medicine). Originally, P3-X63-Ag8 cells were used but NSO-BCL2 cells were used to improve the efficiency of generating hybridomas that produced antibodies against CASPR2. For maintaining fusion partner hybridoma lines P3-X63-Ag8 and NSO-BCL2, cells were grown in complete medium and fed every 2-3 days and split every other feed at 1:5 dilution. For NSO-BCL2, 1.5 mg/mL of G418 (ThermoFisher Scientific, Cat No 10131027) was added to complete medium to select against hybridomas that lose the BCL2 gene. Freshly cultured myeloma cells were centrifuged and washed with cold DMEM to rinse off fetal calf serum (FCS). The rinse was repeated twice. Both splenocytes and myeloma cells were counted to determine the number of live cells. Generally, a normal spleen from the immunized mice generated ∼1×10^8^ cells. Only half of splenocytes were used and the other half were frozen in a mixture of 90% FCS and 10% DMSO. Cells were combined in a 3 splenocytes:1 myeloma ratio. The splenocyte and myeloma mixture was centrifuged to remove all DMEM in preparation of the chemical fusion process. The pellet was loosened by flicking and 0.5 mL of 50% PEG 4000 (pH 7.5–8.0) was slowly added to the pellet over the course of 1 min, while the tube was gently swirled with each drop. After PEG was completely added, the tube was gently swirled in a 37^°^C water bath for 90 sec. To stop the reaction, the PEG mixture was diluted by slowly adding 10 mL of the following solution (4.0 g NaCl, 0.2 g KCl, 0.7 g Na_2_HPO_4_, 0.3 g NaH_2_PO_4_, 1.0 g glucose in 500 mL of dH_2_O) while swirling. Another 10mL of solution was added bringing the total volume to 20 mL The mixture was left standing for 5–10 min, centrifuged and resuspended via drop-and-swirl method in a calculated volume of complete HAT medium (concentration of 3×10^5^ total cells/mL), and plated (100 μL per well) into 96-well plates. These plates were placed in 10% CO_2_, 37^°^C incubator and left undisturbed. Seven days after fusion, wells were fed with 100 μL of warm HAT medium. Hybridomas were fed with medium containing HAT for 3 weeks after fusing or until cloning in agarose and then fed with medium containing HT for 10–14 days to slowly wean them off the selective medium agents. After that time, they were fed with regular medium with 20% FCS. As the positive 24-well plates grow more vigorously, FCS concentration in the medium were lowered to 15% for a week, and then to 10%

### Caspr2-binding ELISA

Supernatant from wells in the 96-well plates with semiconfluent growth of hybridoma cells were assayed by ELISA to determine which wells have anti-CASPR2 antibody producing hybridomas. For simplicity, we used the same name for the hybridoma cell line and the antibody it produced. For ELISA, Costar (Cat No. 3690) half area, 96-well plates were coated with 5 μg/mL of human extracellular portion of CASPR2 protein overnight at 4^°^C. Plates were washed with 1x PBS to remove excess CASPR2 antigen before blocking with 3% Fetal Bovine Serum (FBS) in 1x PBS for 1 h at 37^°^C. Plates were flicked to remove blocking buffer prior to incubation with hybridoma supernatant for 1 h at 37^°^C. Plates were then washed 6x with 1x PBS-0.1% Tween-20 to remove non-binding primary antibody before incubation with the secondary antibody, a 1:1000 dilution of Goat Anti-Mouse IgG(H+L)-AP (Southern Biotech, Cat No. 1036-04). Plates were washed 6x with 1x PBS-0.1% Tween-20 and then incubated with developing solution (AP substrate (Sigma-Aldrich, Cat No S0942), in 0.001 M MgCl_2_ and 0.05 M NaHCO_3_). Plates were incubated at 37^°^C and read at multiple time points (30 min, 45 min, and 1 h) at a wavelength of 405 nM on a 1430 Multilabel Counter Spectrometer (PerkinElmer). Each sample was assayed in duplicate. Each sample that had an average absorbance value greater than the negative control was considered positive and the cells expanded to 24-well plate for future cloning.

### Cloning of hybridomas in soft agarose

Hybridomas producing anti-Caspr2 antibody were cloned. Prior to making agarose plates, the agarose (SeqPlaque™ agarose, Lonza Bioscience, Cat No 50101) was titrated for each lot purchase. Generally, 2.5–3.5 g agarose per 50 mL of dH2O was used. The agarose mixture was dissolved completely before being autoclaved for 15 min on liquid cycle. This agarose stock could be boiled or microwaved to re-melt 3–4 times for cloning. The amount of agarose stock used per 100 mL of medium averaged 6-7.5 mL The goal was for the agarose layer to be loose enough that the clones shake slightly when seen under a microscope but firm enough that colonies do not float across the plate. Three agarose plates were made for each sample. Each plate contained 4-5 mL of cloning medium with agarose for the underlayer and 1 mL for the cell layer. Cloning medium was made from 20% FCS, 10% NCTC-109, 1% nonessential amino acids, 1% pen/strep, 2% HT (50X), and *quantum satis* DMEM with high glucose. For the agarose underlayer, 10% of J774.2 supernatant was added. The cloning medium was warmed to 37^°^C before adding the liquid agarose stock (6-7.5 mL agarose per 100 mL medium). While warm, the cloning medium with agarose was pipetted into each 60-mm petri dish for the underlayer. Dishes were placed on a leveling table and left at 4^°^C, in the cold room for 15 min. To check if the underlayer was solid enough, the dishes were tilted slightly to check for slumping. If there was too much slumping, the plates were maintained in the cold room for another 10 min. If the plates are still slumping after then, the process was started over. While the underlayer was setting, the cell layer was prepped by adding 3 mL of cloning medium with agarose to a 15-mL Falcon tube for each cell line. To create a single cell suspension (around 1,000-2,000 cells per dish) 15 μL of cells was added to 3 mL of cloning medium and agarose. The mixture (1 mL) was added to each dish, drop by drop in a snail spiral pattern so the cells were evenly spaced on top of the agarose underlayer. Dishes were placed on the leveling table at 4^°^C for 15 min. Then, plates were placed in the 37^°^C incubator, 8% CO_2_ for 1 week. Colonies were picked once visible by eye, usually 7–10 days after plating. Before picking colonies, 96-well plates were preconditioned by incubating 100 μL cloning medium per well in 30 wells/cell line for 3 h or overnight. The medium was replaced with fresh cloning medium prior to picking colonies. Colonies were picked via pipette and then placed into a 96-well plate. Once colonies were picked, they were placed in the 37^°^C incubator, 8% CO_2_ for approximately 1 week, when colonies were half confluent, 100 μL of fresh cloning medium was added. Supernatant was collected and screened by ELISA. Colonies that secreted antibody to Caspr2 were transferred into a 24-well plate, expanded and frozen.

### Hybridoma cell culture maintenance

After cloning and confirmation of anti-CASPR2 antibody secretion by ELISA, positive hybridomas were expanded to Fisherbrand™ Surface Treated Sterile Tissue Culture Flasks, Vented Cap (Fisher Scientific, Cat No FB012939) and were transitioned to grow in Gibco™ Hybridoma-SFM medium (ThermoFisher Scientific, Cat No 12045076) following a stepwise process (i.e. 75% complete medium/25% Hybridoma-SFM for a week, 50% complete medium/50% Hybridoma-SFM for a week, and 25% complete medium/75% Hybridoma-SFM for week). Cells were fed every 2–3 days and split every other feed at a 1:5 dilution.

### Antibody purification

Supernatant from hybridoma cultures was collected, centrifuged at 200xg for 5 min (to pellet out cellular debris), and added to Pierce™ Protein G agarose (Thermo Fisher, Cat No. 20399) that had been washed with 1x PBS. Supernatant with agarose was incubated on a rotator overnight at 4^°^C. The next day, the mixture was centrifuged at 1000 rcf for 10 min to pellet out the agarose, which was added to an empty gravity flow column that has been prewashed with 1x PBS. The agarose was washed twice with 1x PBS. Anti-CASPR2 antibody was eluted from agarose using 0.1 M glycine (pH 3.0) into Eppendorf tubes that were preloaded with 1 M Tris-HCl to neutralize the glycine to pH 7.0. The elution fractions with IgG were pooled and dialyzed overnight at 4 ^°^C using a Slide-A-Lyzer Dialysis Cassette (Thermo Fisher, Cat No. A52971) to exchange the antibody buffer to 1x PBS. Antibody concentration was measured using Nanodrop Spectrophotometer and then stored at -20^°^C.

### Sequencing of heavy and light chains of hybridomas

RNA from hybridoma cells was extracted using Qiagen RNeasy Kit (Qiagen) following manufacturer’s instructions. cDNA was generated from extracted RNA using iScript™ cDNA Synthesis Kit (Bio-Rad) following manufacturer’s instructions. The cDNA served as the PCR template for cloning heavy and light chain sequences and follow the method described in[]. x []. A 1% agarose DNA gel was used to confirm the presence of PCR product and the band of the right size was excised and cleaned using QIAquick PCR Purification Kit (Qiagen). The PCR product was then ligated into TOPO-TA vector (Invitrogen) following the manufacturer’s instructions and then transfected into One Shot™ TOP10 Chemically Competent *E. coli* bacteria (Invitrogen). Bacteria were plated on ampicillin LB agar plates and placed upside down in 37°C incubator overnight. The next day colonies were picked and sent to Azenta Life Sciences for Sanger sequencing. The sequencing results were then aligned against the IMGT® database to determine specific heavy and light chains.

### IgG binding ELISA

Costar (Cat No. 3690) half area, 96-well plates were coated with 10 μg/mL of unlabeled anti-mouse IgG, IgG1, IgG2b, IgG2c, IgG3 overnight at 4^°^C. Plates were blocked with 3% FCS for 1 h at 37°C. Hybridoma-generated anti-Caspr2 antibodies were tested against each isotype by adding supernatant to the ELISA plate followed by an AP conjugated antibody to the same isotype (e.g. anti-mouse IgG1 AP was used as a secondary antibody for a well that was coated with anti-mouse IgG1). Plates were incubated with developing solution (AP substrate, Sigma-Aldrich, Cat No S0942, in 0.001 M MgCl_2_ and 0.05 M NaHCO_3_), incubated at 37^°^C, and read at 30 min at a wavelength of 405 nM on a 1430 Multilabel Counter Spectrometer (PerkinElmer).

### Characterization of IgG binding by ELISA

Costar (Cat No. 3690) half area, 96-well plates were coated with 5 μg/mL of human extracellular portion of CASPR2 protein overnight at 4^°^C. Plates were washed with 1x PBS to remove excess CASPR2 antigen before blocking with 3% FBS in 1x PBS for 1 h at 37^°^C. Plates were flicked to remove blocking buffer prior to incubation with 10 μg/mL of anti-CASPR2 antibody, diluted in 0.3% FBS for 1 h at 37^°^C. For comparing binding affinities, a serial dilution of anti-Caspr2 antibody concentration was used (in μg/mL: 40, 20, 10, 5, 2.5, 1.25, 0.75, 0.375). Plates were washed 6x with 1x PBS-0.1% Tween-20 to remove primary antibody before incubation with the secondary antibody, a 1:1000 dilution of Goat Anti-Mouse IgG(H+L)-AP (Southern Biotech, Cat No. 1036-04). Plates were washed 6x with 1x PBS-0.1% Tween-20 and then incubated with developing solution (AP substrate (Sigma-Aldrich, Cat No S0942), in 0.001M MgCl_2_ and 0.05M NaHCO_3_). Plates were incubated at 37^°^C and read at 30 min at a wavelength of 405 nM on a 1430 Multilabel Counter Spectrometer (PerkinElmer).

### Caspr2 binding cell based assay

HEK cells (5,000,000) were seeded in 100 × 20 mm Falcon® Tissue Culture-treated dish (Fisher Scientific, Cat no 353003) for each transfection. At 24 h after plating, cells were transfected via Fugene (Promega, Cat no #E269A) with human Caspr2 GFP construct, mouse Caspr2 GFP construct, or nothing. At 48–72 h after transfection, the live cell-based assay was performed. Cells were lifted using a mixture of 5% FBS (R&D Systems, Cat no #S11150H) with 1mM EDTA in HBSS (Gibco, Cat No #14175, 095), scraped with a cell scrapper (Corning, Cat No 353085) and collected into a 50-mL conical tube. Cells were spun down at 1200 rpm for 5 min at 4°C. The pellet was resuspended in 5% FBS, 1mM EDTA in HBSS before counting on a hemacytometer. Cells (500,000 per well) were transferred into a non-treated 96-well round bottom plate (Falcon, Cat No 351177) and used to test each anti-Caspr2 antibody for binding. Cells were spun down in the plate at 1500 rpm (5 min at 4°C) and incubated with 10 μg/mL of primary antibody (anti-CASPR2 antibody or their respective isotype control) or 1:400 dilution of serum from a mouse immunized with the extracellular portion of the human CASPR2 as a positive control and fluorescent viability dye (FVD 506, Invitrogen eBioscience, Cat No 65-0866-14) for 1 h on ice, protected from light. Cells were washed with 5% FBS with 1mM ETDA in HBSS before adding secondary antibody, 1:400 Alexa Fluor 594 goat anti-mouse IgG (Invitrogen, Cat No A11032) for 30 min on ice, protected from light. Cells were washed in 5% FBS with 1mM EDTA in HBSS and fixed in 1% PFA (diluted from 32% PFA, Electron Microscopy Sciences, Cat No 15714-S) diluted in 5% FBS with 1mM EDTA in HBSS. Live cells were gated on FVD 506.

#### Timed pregnancies via trio breeding

### Generation of adult mouse brain sections

Male C57BL/6 mice (12–20 weeks old) underwent transcardial perfusion following standard protocols. For PFA fixed sections, mice were first perfused with pre-perfusion solution (see appendix for recipe) followed by 4% PFA diluted in 0.2 M PB. Brains were carefully removed from the skull and put into vials containing 20 mL of 4% PFA solution and post-fixed for 2 h. Brains were transferred to 30% sucrose solution for 24–48 h before freezing in Tissue-Tek® OCT compound (Sakura). Brains were stored in -80°C until sectioning. For snap frozen brain sections, mice were perfused with pre-perfusion solution prior to brain removal, brains were then snap frozen in -40°C methylbutane and stored at -80°C until sectioning. For both PFA-fixed and snap frozen brains, 14-μm sections were cut on a cryostat and mounted onto Fisherbrand Superfrost Plus Microscope Slides (FisherScientific, Cat No 22-037-246). Sections were stored at -80°C until staining.

### Generation of E15.5 mouse sections

Timed pregnancies were generated with trio breeding, in which a male was cohoused with 2 females overnight (14 h) and separated the following morning (day E0.5). Mice were checked for pregnancy on day E12.5 by weight, in which an increase of 2 g (or more) and a rounding of the belly was taken as a sign of pregnancy.

Dams were euthanized using CO_2_ following IACUC guidelines on day E15.5. Embryos were extracted via c-section and were washed in ice cold Gibco™ HBSS (ThermoFisher Scientific, Cat No 14175095) prior to fixation in 4% PFA, 4% sucrose for 4 h at room temperature. Embryos were incubated in a sucrose gradient (10% sucrose for 45 min at 4°C, 20% sucrose for 2 h at 4°C, 30% sucrose overnight at 4°C) prior to freezing in Tissue-Tek® OCT compound (Sakura). Full embryo, 12-μm sagittal sections were cut on the cryostat and mounted onto Fisherbrand Superfrost Plus Microscope Slides (FisherScientific, Cat No 22-037-246). Sections were stored at -80°C until staining.

### Generation of mouse primary neuronal cultures

Postnatal day 0 (P0) to postnatal day 3 (P3) pups were used to generate primary neuronal cultures; 6–8 pups were euthanized by decapitation via large sharp scissors. The cortex was isolated by dissection and all cortices were pooled together in a solution of HDGH on ice. The cortices were transferred into a sterile 60-mm petri dish filled with 2mL of PPD. The tissue was mechanically cut using a sterile Pasteur pipette and transferred into a 15-mL conical tube. PPD (3 mL) was added to the petri dish to catch any remaining tissue and then pooled with previous 2 mL of tissue in the conical tube. Each conical tube was incubated at 37°C for a total of 15 min with gentle rocking every 5 min. The tissue was mechanically dissociated using a fire polished, glass pipette by gentle trituration 15 times. The tissue was incubated at 37°C for 15 min with gentle rocking every 5 min as before. The tissue was subjected to another round of mechanical dissociation using another fire polished, glass pipette with an even narrower opening than the one used previously. Cells were centrifuged at 250xg for 5 min at 4°C. The supernatant was removed and the cellular pellet was resuspended with 5 mL of warm NBGxPPS. Cells were filtered through a 0.4-μm filter prior to counting using a hemacytometer. For immunohistology, 200,000 cells per well were seeded onto glass coverslips (Electron Microscopy Sciences Circular Cover Glass #1.5, 18mm, Fisher Scientific Cat No 50-948-975) coated with 1x poly-D-Lysine. For capillary western, 1,500,000 cells per well were seeded onto Corning BioCoat Poly-L-Ornithine/Laminin Multiwell Plates (Corning, Cat No 354658). The next morning, the plates were vigorously shaken prior to a medium change to remove dead cells and debris. Cells were fed every 3 days until day of experiment.

### Immunohistochemistry on PFA fixed adult brain sections

Sections were rehydrated in 1x PBS for 5 min twice prior to antigen retrieval in 10 mM sodium citrate buffer at 95°C for 2 min. Sections were washed in 1x PBS for 5 min and blocked in 3% BSA in 0.1% Triton X-100 at room temperature for 1 h, incubated with 10 μg/mL of hybridoma generated anti-CASPR2 antibodies overnight at 4°C, washed with 1x PBS-0.1%Tween-20, and then incubated with secondary antibody anti-mouse Alexa Fluor 488 (ThermoFisher Scientific, Cat No A32723, 1:400) for 1 h at room temperature, protected from light. Sections were then stained with 0.5 μg/mL DAPI for 5 min prior to being coverslipped with DAKO mounting medium (Agilent, Cat No S3023). Microscopy images were taken at 20x.

### Immunohistochemistry on snap frozen adult brain sections

Sections were fixed in 95% EtOH in -20°C for 30 min, followed by 1 min in acetone at -20°C. Sections were blocked in 3% BSA in 1x PBS-0.1% Triton X-100 at room temperature for 1 h, incubated with 10 μg/mL of hybridoma generated anti-Caspr2 antibodies overnight at 4°C, washed with 1x PBS-0.1% Tween-20, and incubated with secondary antibody anti-mouse Alexa Fluor 488 (ThermoFisher Scientific, Cat No A32723, 1:400) for 1 h at room temperature, protected from light. Sections were stained with 0.5 μg/mL DAPI for 5 min prior to being coverslipped with DAKO mounting medium (Agilent, Cat No S3023). Microscopy images were taken at 20x.

### Immunohistochemistry on E15.5 sections

Sections were rehydrated in 1x PBS for 5 min twice prior to antigen retrieval in 10 mM sodium citrate buffer (see appendix for recipe) at 95°C for 2 min. Sections were washed in 1x PBS for 5 min and blocked in blocking buffer (3% BSA, 3% FCS in 0.1% Triton X-100) at room temperature for 1 h prior to incubation with 10 μg/mL of hybridoma generated anti-CASPR2 antibodies conjugated to Alexa Fluor 488 overnight at 4°C. Antibodies were labelled using Alexa Fluor 488 Antibody Labeling Kit (ThermoFisher Scientific, Cat No A20181). Sections were washed with 1x PBS-0.1% Tween-20 and stained with 0.5 μg/mL DAPI for 5 min prior to being coverslipped with DAKO mounting medium (Agilent, Cat No S3023). Microscopy images were taken at 20x.

### Immunohistochemistry on mouse primary neuronal cultures

DIV14 neuronal cultures were fixed in 4% sucrose, 4% PFA at room temperature for 10 min. Cells were blocked with 4% BSA in 1x PBS for 30 minutes at room temperature. Cells were incubated in primary antibody; 10 μg/mL of anti-Caspr2 (P11G7, P9C7, P6B2, or P13F4) and a commercial anti-CASPR2 antibody (EMD Millipore, Cat No ABN1380, 1:1000) diluted in blocking buffer overnight at 4°C. Cells were incubated in secondary antibodies; anti-mouse Alexa Fluor 488 (ThermoFisher Scientific, Cat No A32723, 1:400) and anti-rabbit Alexa Fluor 594 (ThermoFisher Scientific, Cat No A32740, 1:400) diluted in blocking buffer for 1 h at room temperature, protected from light. Cells were incubated with DAPI (ThermoFisher Scientific, Cat No 62248, 0.5 μg/mL) 5 min at room temperature, protected from light. The glass coverslips were then mounted onto Fisherbrand Superfrost Plus Microscope Slides (FisherScientific, Cat No 22-037-246) using DAKO mounting medium (Agilent, Cat No S3023), sealed with clear nail polish, and then visualized by microscopy. Microscopy images were taken at 20x on Zeiss Brightfield Microscope using a Colibri LED system.

### Generation of behavioral cohorts

Timed pregnancies were set up by trio breeding as described above. At E13.5, 200 μg of anti-Caspr2 antibody (P11G7, P9C7, P6B2, or P13F4) or an isotype control antibody was injected into pregnant dam via retroorbital injection. Pregnant dams were separated and single housed. At least 5 litters with each antibody treatment were generated so at least 10 mice were in each treatment group segregated by gender. After weaning, mice were transferred to the reverse light cycle room for behavioral testing.

### Behavioral assessments

Mice exposed to anti-CASPR2 antibodies *in utero* were housed under a reversed dark (9:00–21:00) and light (21:00–9:00) cycle, with *ad libitum* access to food and water. All manipulations were conducted during the dark phase, at least 1 h after turning the lights off. Mice were handled 3x (15 min sessions on separate days) before behavioral assessments, which were performed in the following order: light-dark-chamber task, open-field task, and social interaction task. During experimentation, the investigator was blinded to the antibody exposure status of the mice. All the tasks were recorded with a centrally placed video camera directly above each arena which fed the signal to the tracking software (EthoVision XT 14.0, Noldus, Attleboro, MA, USA).

### Light-dark chamber task

This test examined anxiety by measuring the mice’s natural aversion to bright light. Each mouse was placed for 10 min in the Noldus light-dark box (Noldus, Attleboro, MA, USA),which consisted of a chamber with a large open compartment (which was highly illuminated) and a small dark compartment. The Noldus light-dark box is made from IR translucent materials, which in combination with the IR backlight and IR-sensitive camera, makes the apparatus suited for video tracking with EthoVision software. The apparatus was cleaned prior to each trial with 70% ethanol followed by water and wiped dry.

### Open field task

This test examined locomotor activity and anxiety-like behavior by placing the mice in the center of a square arena (40 × 40 cm) with gray walls (35 cm high) and allowing them to freely explore the chamber during two sessions (15 min each) separated by ∼2 h. The sessions were recorded with a centrally placed video camera directly above the arena which fed the signal to EthoVision software, which was used for automated analysis of animal behaviors including distance traveled, velocity, time spent moving, time spent in the center of the arena (20 × 20 cm^2^). The arena was cleaned in between each trial with 70% ethanol followed by water and wiped dry.

### Social interaction task

Prior to beginning the task, the “stimulus” mice were lightly anesthetized, shaved, and their backs were colored with a non-toxic marking ink, allowing these mice to be differentiated from the experimental mice. Moreover, each stimulus-mouse was placed into a chamber (30 × 30 cm, 40 cm high) for 20 min to familiarize itself with the environment. For the test proper, the stimulus mouse was introduced to the chamber together with an experimental mouse and the two animals were allowed to interact for 10 min. The social interaction module of the EthoVision software was used to detect body contact, based on the body contours of the two animals.

### Golgi-Cox staining and Sholl analysis

Brains were prepared for morphometric analysis, as previously described (46), with Rapid GolgiStain™ Kit (FD Neuro Technologies), a silver staining method that permits visualization of entire neurons. Comparable sections across all animals were imaged on an Axioimager Z1 microscope (N.A.=0.75, z-step=5µm). There were 3-5 animals in each group, and for each animal there were at least 7 neurons traced and measured using Neurolucida® (MBF Bioscience). Dendritic length measurements were compiled according to a Sholl analysis (6) (Stereoinvestigator/Neurolucida®) for a linear mixed model analysis (R program v.3.6.1 with Rstudio) to test for group differences and generate interclass correlation values.

### Anti-Caspr2 antibody target domain identification

HEK293T cells were transfected with human full-length Caspr2 cDNA or mutant constructs, each missing one of the domains. These mutant constructs were generated as described previously (44). Reactivity against these constructs was then tested through cell-based assay with live-staining using established methods [45]. In brief, 48 h after transfection, antibodies were diluted to 5 µg/ml in growth medium (DMEM+Glutamax, 10% FBS, 1% Streptomycin/Penicillin, 1% MEM non-essential amino acid solution) and added to the cells. After 1 h incubation (37ºC), medium was removed and cells were fixed with 4% PFA. After washing, Alexa Fluor 488 coupled goat anti-human antibody or Alexa Fluor 488 coupled goat anti-mouse IgG (1:1000; Dianova CAT#:109-545-003 1:1000; Invitrogen CAT#: A-11029) was added and left to incubate for 2 h at room temperature. Images were then obtained using widefield imaging on a Leica SPE. Reactivity against the discoidin domain was confirmed through a CBA using an additionally generated construct containing only the discoidin domain.

## ACKNOWLEDGEMENTS

We thank Joe Gallagher for help with behavioral assays. Funding was provided by the National Institute of Health (NIH) grant 5P01AI073693 (to BD). PH acknowledges the Department of Defense (DOD) impact award W81XWH1910759. PR received funding from the German Research Foundation (DFG) grants FOR3004, PR1274/9-1, clinical research unit 5023/1 ‘BECAUSE-Y’ (project number 504745852), the Helmholtz Association (HIL-A03 BaoBab), and the German Federal Ministry of Education and Research (Connect-Generate 16GW0279K). JK was supported by the Berlin Institute of Health at Charité Clinician Scientist program and received funding from German Research Foundation (DFG) grant FOR3004 (project number 415914819).

## Notes

### Competing Interest Statement

The authors have declared no competing interest.

